# Opportune cell culture conditions permit transfer of the neurosphere paradigm to cells of the cerebellar granule cell lineage by being permissive to sonic hedgehog signaling

**DOI:** 10.1101/096859

**Authors:** Constantin Heil

## Abstract

The *in vitro* study of neural progenitors is based around the idea that defined culture conditions can emulate an environment which maintains an undifferentiated state and self-renewal capability of explanted progenitor cells, culminating in the growth of neurospheres. Neurosphere culture systems are used to study cells of physiological origin as well as transformed cells. In the case of the cerebellar granule cell lineage, neurosphere cultures have been produced from medulloblastoma tumors, while non-transformed granule progenitors are reported to be transient *in vitro.* SHH signaling is a key signaling pathway with roles in morphogenesis and cell proliferation in the central nervous system, in particular, involved in mitogenic signaling within the context of postnatal cerebellar granule cell expansion. In this paper, I demonstrate that in addition to commonly used mitogens such as EGF and bFGF, the SHH pathway agonist SAG, as well as genetic activation of SHH signaling by PTCH1 deletion, can lead to the growth of neurospheres from postnatal day 7 (p7) cerebellar explants.

I also show that these neurosphere adhere to the cerebellar GCP lineage.. Strikingly, mSS cells can be maintained indefinitely in culture, as demonstrated by extensive clonogenic capability, assayed over a period of 10 weeks. In the context of self-renewal, I assay gene expression of POU3f2, POU5f1, NANOG, and SOX2, genes associated with neural progenitors, which I find expressed in mSS cells. Importantly, mSS cultures are continuously dependent on SAG for their clonogenic potential and SHH pathway activation, assayed by expression of GLI1, PTCH1 and NMYC. In addition to the aforementioned extensive self-renewal capability, mSS neurosphere cultures also maintain the ability to differentiate. *In vitro* differentiation leads to formation of cells with typical granule cell morphology and which are positive for beta3-tubulin and express GABRA6.

Overall, my work demonstrates that by applying culture conditions which are tailored towards biological characteristics of specific regions of the central nervous system the paradigm of the neurosphere can be expanded to include lineages not previously studied in this way. In particular, I apply this principle to unmask the property of cells from the GCP lineage as having extensive self-renewal capability *in vitro.*

## Introduction

Development of the mammalian cerebellum continues after the birth of the animals, particularly manifesting itself in the form of a transitory layer of proliferating cells on the surface of the developing cerebellum, i.e. the external granule layer (EGL). Cells within the EGL undergo cell division, migration and finally terminally differentiate to give rise to the internal granule layer (IGL). The completed IGL houses the population of mature cerebellar granule neurons, which are glutamatergic neurons that are the most abundant cells in the mammalian body.

Proliferation of cerebellar granule progenitors (GCPs) is transient and is finely regulated by a plethora of extracellular signals, of which sonic hedgehog (SHH) signaling has been demonstrated as being necessary and sufficient to drive GCP expansion (Wechsler-Reya and Scott, 1999). However other signals have been demonstrated to either potentiate this response, such as IGF-1 (Browd et al., 2006), or to contribute to the attenuation of the SHH dependent mitogenic response (Rios et al., 2004). During development of the cerebellum, proliferation of GCPs eventually ceases at approximately 5 months *post-partum* in humans and in mice approximately 21 days after birth. Prolonged proliferation due to either enhancement of mitogenic signaling or disruption of active inhibitory cues has been linked to the genesis of type 2 medulloblastoma (Northcott et al., 2012).

The transient proliferation of GCPs has been extensively studied *in vitro* and the reported methods of culture of these cells lead to the production of primary cultures, which recapitulate the limited proliferative capability of GCP lineage cells as also displayed during normal *post-natal* development (Miyazawa et al., 2000; Wechsler-Reya and Scott, 1999). The observation that GCPs cease proliferation and differentiate in primary cultures despite even prolonged treatment with exogenous SHH protein or agonists of the SHH pathway has led to the hypothesis that GCPs posses intrinsic mechanisms of regulation of cell cycle exit and differentiation (Kessler et al., 2009).

In addition to being cultured as adherent cells, neural progenitors can also be cultured in defined media without serum where they exist as floating, proliferating cell aggregates known as neurospheres. Neurosphere cultures were pioneered in the 1990s (Reynolds and Weiss, 1992) and are often used as models of neural stem cells. This is due to their properties, which match the operative definition of a neural stem cell, these being multi-potency and extensive self-renewal capabilities (Pastrana et al., 2011; Reynolds and Weiss, 1992).

Neural progenitors that undergo rounds of clonal expansion followed by differentiation are known as transit amplifying (TA) progenitor cells (Butts et al., 2014). In addition to GCPs, TA cells have been identified in other systems of *post-natal* neural development such as the dentate gyrus of the hippocampus and the adult sub-ventricular zone (SVZ) (Capdevila et al., 2016; Gonçalves et al., 2016).

Interestingly, *in vivo* studies have cast doubt on the *in vivo* stem cell identity of *in vitro* cultured neurospheres (Pastrana et al., 2011). This is best exemplifed by Doetsch et. al. in which the authors develop a labeling strategy to retrospectively identify the origin of SVZ derived neurospheres as being not of the stem cell compartment but of the TA compartment (Doetsch et al., 2002).

The cellular hierarchies which determine the dynamics of neural tissue are also relevant in the context of cancer biology. In that context of brain tumors, it has been reported that tumor initiating cells or tumor maintaining cells posses properties of immature neural progenitors (Ahlfeld et al., 2013; Capdevila et al., 2016; Vanner et al., 2014). Further, diverse methods have been proposed for the culture of brain tumor derived cells (Rahman et al., 2015). Choice of opportune conditions can lead to more or less relevant cell lines (Sasai et al., 2006).

In this work I explore alternate culture conditions for GCP lineage cells. I begin with characterization of cell cultures obtained from murine medulloblastoma and extend these studies to non-transformed GCPs. Importantly, I demonstrate that GCPs can be cultured as neurospheres solely under conditions which permit activity of the SHH signaling pathway. Further, I discover that these cell cultures can be maintained for extensive periods, remaining dependent on SHH signaling. Additionally, under these conditions, GCP cells maintain the potential to differentiate into post-mitotic granule cells.

## Materials and Methods

### Establishment of cultures from murine central nervous system

Postnatal day 7 (p7) mice were sacrificed by decapitation and tissues were collected in HBSS supplemented with 0.5% glucose and penicillin-streptomycin, grossly triturated with serological pipette and treated with DNAse I to a final concentration of 0.04% for 20 min. Finally, cell aggregates were mechanically dissociated using pipettes of decreasing bore size to obtain a single-cell suspension. SCs were cultured as neurospheres in selective medium after centrifugation, DMEM/F12 supplemented with 0.6% glucose, 25 μg/ml insulin, 60 μg/ml N-acetyl-L-cystein, 2 μg/ml heparin, 20 ng/ml EGF, 20 ng/ml bFGF (Peprotech, Rocky Hill, NJ) or 1μM SAG (Adipogene), 1X penicillin-streptomycin and B27 supplement without vitamin A. SAG dosage was 1μM unless otherwise specified.

For differentiation, cells were mechanically dissociated and plated on polylysinated cell culture dishes in the following medium: Neurobasal (Invitrogen) supplemented with glutamine, penicillin-streptomycin, 0.5% fetal bovine serum and B27 supplement.

### Clonogenic assay

Cells were pelleted and dissociated by incubation with Accutase (Sigma Aldrich) in concomitance with continuous pipetting to obtain a single cell suspension. Cells were counted with a hemocytometer and were diluted to obtain a suspension concentrated 0.267 cells/μl. Cell suspension was distributed amongst a 96 well plate, pipetting 75μl into each well.

Respective treatments were prepared with double the final concentration and 75 μl were added to each well for a final volume of 150μl concentrated 0.134 cells/μl, for a total of 20 cells per well. After 3 days medium was replenished and an additional 3 days later the number of spheres per well was recorded. Exhibited data represents averages of 2 independent experiments, each having 24 wells per treatment point.

### RNA preparation and quantitative reverse transcription-PCR

Total RNA extraction was carried out with TRIzol reagent (Invitrogen). For quantitative reverse transcription-PCR (QPCR), total RNA (1μg) was reverse transcribed using the Gene Amp kit (Applied Biosystems) and subjected to PCR amplification using SYBR Green PCR Master Mix (Applied Biosystems) or TaqMan PCR Master Mix (Applied Biosystems). Samples underwent 40 amplification cycles (95C for 10 seconds; 60C for 1 minute) in a ABI Prism 7900 machine (Applied Biosystems). TaqMan probes (Applied Biosystems) were used to detect HPRT, GLI1, PTCH1, GABRA2 and MYCN. For SYBR green QPCR the following primers were used:

Nestin FW GCAGGCCACTGAAAAGTTCC
Nestin RV CACCTTCCAGGATCTGAGCG
SOX2 FW CAGGAGTTGTCAAGGCAGAGA
SOX2 RV CTTAAGCCTCGGGCTCCAAA
Nanog FW CGGTGGCAGAAAAACCAGTG
Nanog RV GGTGCTGAGCCCTTCTGAAT
POU3F2 FW CCTTTAACCAGAGCGCCCA
POU3F2 RV AGGCTGTAGTGGTTAGACGC
POU5F1 FW GGCTTCAGACTTCGCCTTCT
POU5F1 RV TGGAAGCTTAGCCAGGTTCG
ATOH1 FW CCCGTCAAAGTACGGGAACA
ATOH1 RV CTCGTCCACTACAACCCCAC

The heatmap in Figure 1a was made in R using the package *heatmap.2* by Gregory R. Warnes. Z-scores of ΔΔct values are graphed.

**Figure 1.**
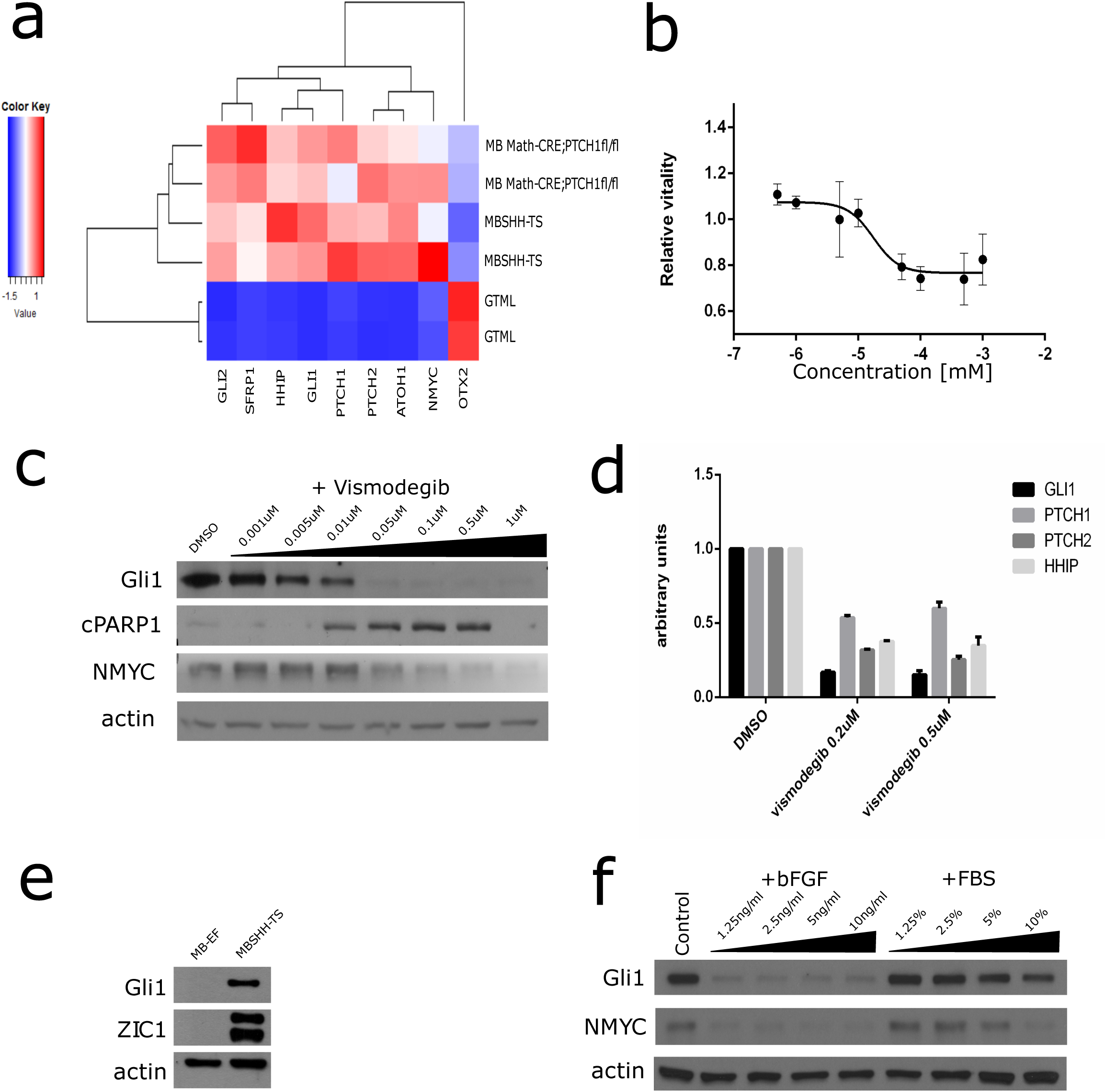
Properties of medulloblastoma cell lines from ATOH1-CRE; PTCH1 fl/fl murine tumors. a) Heatmap summarizing QPCR data comparing MBSHH-TS cell lines with their respective original tumors and with GTML murine medulloblastomas. b) XTT assay measuring vitality of MBSHH-TS cells as function of vismodegib dosage. c) Western blot measuring levels of GLI1, cPARP1 and NMYC in response to different vismodegib dosages. d) QPCR data depicting SHH pathway gene expression levels after vismodegib treatment. e) Comparison of GLI1 and ZIC1 protein levels between MB-EF and MBSHH-TS cells originating from same initial tumor. f) GLI1 and NMYC protein levels in MBSHH-TS cell after exposure to either bFGF of fetal bovine serum (FBS)

### Protein extraction and Western blot

Total protein extracts were obtained in RIPA buffer [50 mmol/L Tris (pH 8), 150 mmol/L NaCl, 0.5% Sodiumdeoxycolate, 0.1% SDS, 1% NP40, 0.001 mol/L EDTA and a mix of protease inhibitors]. Protein extracts were separated by SDS-PAGE and blotted onto nitrocellulose membrane (PerkinElmer). Membranes were blocked with 5% nonfat dry milk and incubated with primary antibodies (Abs) at the appropriate dilutions. Abs were as follows: goat anti–b-actin, rabbit anti CCND1 and mouse anti-MYCN (Santa Cruz Biotechnology), mouse anti-Gli1, #2643 (Cell Signaling Technology Inc). Immunoreactive bands were visualized by enhanced chemoluminescence (Perkin Elmer).

### Immunofluorescence

Cells were fixed in 4% paraformaldehyde and washed with PBS. Samples were permeabilized in 0.1% triton-X 100 in PBS for 15 minutes at room temperature, washed in PBS and blocked in 5% BSA for 30 minutes at room temperature. Primary antibodies were diluted in 1% BSA, incubated overnight at +4 degrees celsius and were the following:

mouse anti-GFAP monoclonal antibody MAB360 (Millipore)
mouse anti-Neuronal Class IIIb-Tubulin (TUJ1) monoclonal antibody MMS-435P (Covance)

Samples were washed in PBS and incubated in Alexa Fluor 555 Goat anti-Mouse (Life Technologies) together with Hoechst nuclear dye for 20 minutes at room temperature. After further washes in PBS slides were mounted in PBS/glycerol and analyzed under fluorescence microscope. Images were analyzed and scored using ImageJ.

## Results

### Generation of relevant murine tumor spheres from conditional knockout disease model

It was recently reported that murine primary MB explants from the PTCH +/− model could be cultured as tumor spheres which maintain activity of the SHH pathway (Zhao et al., 2015). I extend this approach to another model of murine SHH type MB, the ATOH1-CRE; PTCH1 fl/fl mouse. As reported previously, cells proliferated in the form of spheres. These cells, herein named SHH medulloblastoma tumor spheres (MBSHH-TS) express relevant lineage specific and SHH pathway associated genes and consequently cluster together with their respective primary tumor tissues of origin (**Fig. 1a**). Conversely, MBSHH-TS cell lines are readily distinguishable from the type 4 MB model GTML. MBSHH-TS cells further demonstrated nanomolar sensitivity to the smoothened inhibitor vismodegib, as shown by XTT assay (**Fig. 1b**). To confirm that vismodegib sensitivity is due to inhibition of the SHH pathway, I assayed the expression of NMYC and GLI1 by western blot, both of which decreased in a dose dependent manner (**Fig. 1c**).

Further, cleaved PARP1 levels increased in a dose dependent manner, thus recapitulating type 2 MB behavior in this *in vitro* model. In addition to the aforementioned western blot data I also assayed the expression of *gli1*, *ptch1*, *ptch2* and *hhip* genes, all of which responded negatively to vismodegib treatment (**Fig. 1d**).

To underline the novelty and significance of these types of cell lines, I compare MBSHH-TS cells to cells from the same tumor explant, which were cultured in neurosphere medium supplemented with EGF and bFGF. These cells do give rise to neurospheres, however such cells do not express detectable levels of either GLI1 or ZIC1 protein (**Fig. 1e**), indicating that they may not be relevant towards the study of SHH subtype medulloblastoma.

The finding that growth factors may not support SHH signaling reflects literature which describes that bFGF treatment inhibits the signaling pathway (Emmenegger et al., 2013). Consistent with these observations, treatment of established MBSHH-TS cells with either bFGF or serum led to diminishment of GLI1 protein expression and the SHH pathway downstream effector NMYC (**Fig. 1f**).

### The postnatal cerebellum supports generation of novel neurosphere cultures which express an activated SHH pathway

In the context of the forebrain, the discovery that cells could be cultured as neurospheres eventually led to another discovery, in the form that brain tumors, such as glioblastoma multiforme (GBM), could also lead to formation of such cultures (Vescovi et al., 2006). There exists, therefore, some kind of duality between transformed and non-transformed cells, in which the same culture conditions can lead to generation of different cell lines based on physiological or diseased status of the starting material.

To explore the possibility that this duality could be extended to the context of SHH type MB and non-transformed GCP cells, I reasoned that cells originating from postnatal murine cerebellar explants should be cultured under conditions which support SHH pathway activation. I therefore exposed cell pools originating from dissociated p7 murine cerebella to serum-free neurosphere culture conditions, supplemented with the growth factors bFGF and EGF (EF conditions) and/or SAG, a small molecule agonist of the SHH signaling pathway.

I observed that after 7 days in culture, primary explants treated with EF conditions lost expression of the SHH pathway mediator GLI1 while cells exposed to SAG maintained GLI1 expression (**Fig. 2a**). Further, I observe that GLI1 expression is also undetectable in cells exposed to EF together with SAG, thus confirming previous observations that growth factor signaling can actively inhibit SHH signaling. In addition to measuring GLI1 protein levels under these conditions, I also carried out a clonogenic assay in order to quantify the formation of neurospheres under the aforementioned conditions.

**Figure 2.**
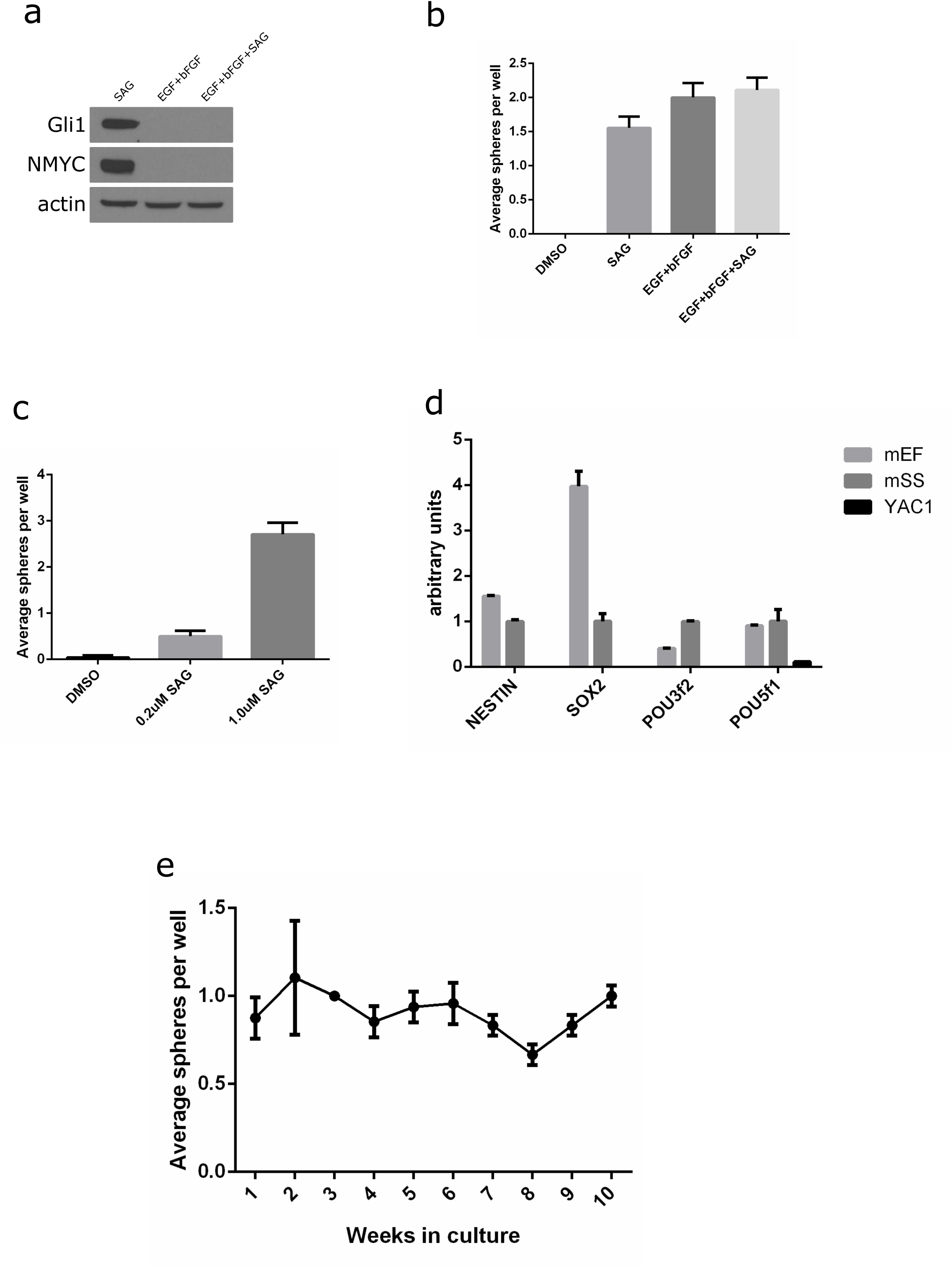
Generation of novel SHH pathway active neurospheres from murine cerebellum. a)Western blot of different primary cultures after 7 days in culture. b) Clonogenicity of explants as in a) and control (DMSO) treated explants. c) Clonogenicity of SAG cultured cells exposed to different concentrations of SAG. d)QPCR of different established cell types and YAC1 mouse lymphoblastoid cells. e) Serial clonogenic assay carried out each week, over a period of 10 weeks, from SAG cultured explants.

I observe formation of spheres under all the experimental conditions tested, while no clonogenicity was observed in control (DMSO) treated explants (**Fig. 2b**). I therefore conclude that while diverse culture conditions can give rise to neurosphere cultures from the developing murine cerebellum, these cultures only express an active SHH pathway when cultured in the presence of SAG without the other tested growth factors.

After having established SHH pathway expressing neurospheres, I challenged cerebellar explants with a lower dose of SAG to investigate sensitivity to SAG levels. Interestingly, 0.2μM SAG led to significantly decreased sphere formation from primary cerebellar cells (**Fig. 2c**).

Neurospheres express markers of neural progenitors and stem cells (Pastrana et al., 2011). To assess whether these novel SAG exposed neurospheres mimic this hallmark I assayed expression of the neural progenitor associated gene *nestin*, as well as the stem-cell factors *sox2* and it’s binding partners *pou5f1* (*oct4*) and *pou3f2*(*brn2*), relative to the murine lymphoblastoid line YAC1. SAG exposed cells expressed all the aforementioned markers, even if *sox2* was expressed at lower levels (**Fig.2d**).

Neurospheres are known for their extensive proliferative capability, while GCP cultures have normally been described as being of transient nature, consistent with their developmental origin. This led me to investigate and, subsequently, challenge the long-term proliferative properties of my novel SAG grown neurospheres by measuring clongenicity for a period of 10 weeks, corresponding to 20 passages. I observe that SAG supports continuous clonogenicity of these cells throughout the duration of the experiment (**Fig. 2e**).

I chose the time frame of 10 weeks for this experiment, however, I report that these cells proliferate indefinitely, as far as I can judge. The extensive proliferation of GCP lineage neurospheres permits the establishment of long-lived cell lines from cerebellar explants, these lines will be referred to as murine SAG induced spheres (mSS) from now on, while neurospheres grown in EF conditions will be referred to as murine EGF/bFGF dependent spheres (mEFS).

### Hedgehog pathway active, cerebellum derived neurospheres belong to the GCP lineage

Next, I sought to investigate the identity of SAG dependent neurospheres. Murine neurospheres have been known to originate from diverse sites along the neuraxis, including the sub-ventricular zone (SVZ). To inquire the specificity of establishment of SAG dependent neurospheres as a function of anatomic location I exposed SVZ and cerebellar explants from p7 brains to either EF conditions or SAG and assayed sphere formation in terms of clonogenicity.

While I replicate the observation that neurospheres are generated from the SVZ and cerebellum in the presence of the two growth factors (Lee et al., 2005), SAG could only lead to the generation of neurospheres from the cerebellum (**Fig. 3a**). This observation led me to hypothesize that the *in vivo* origin on these cells could be a cell type unique to the cerebellum at this developmental stage.

**Figure 3.**
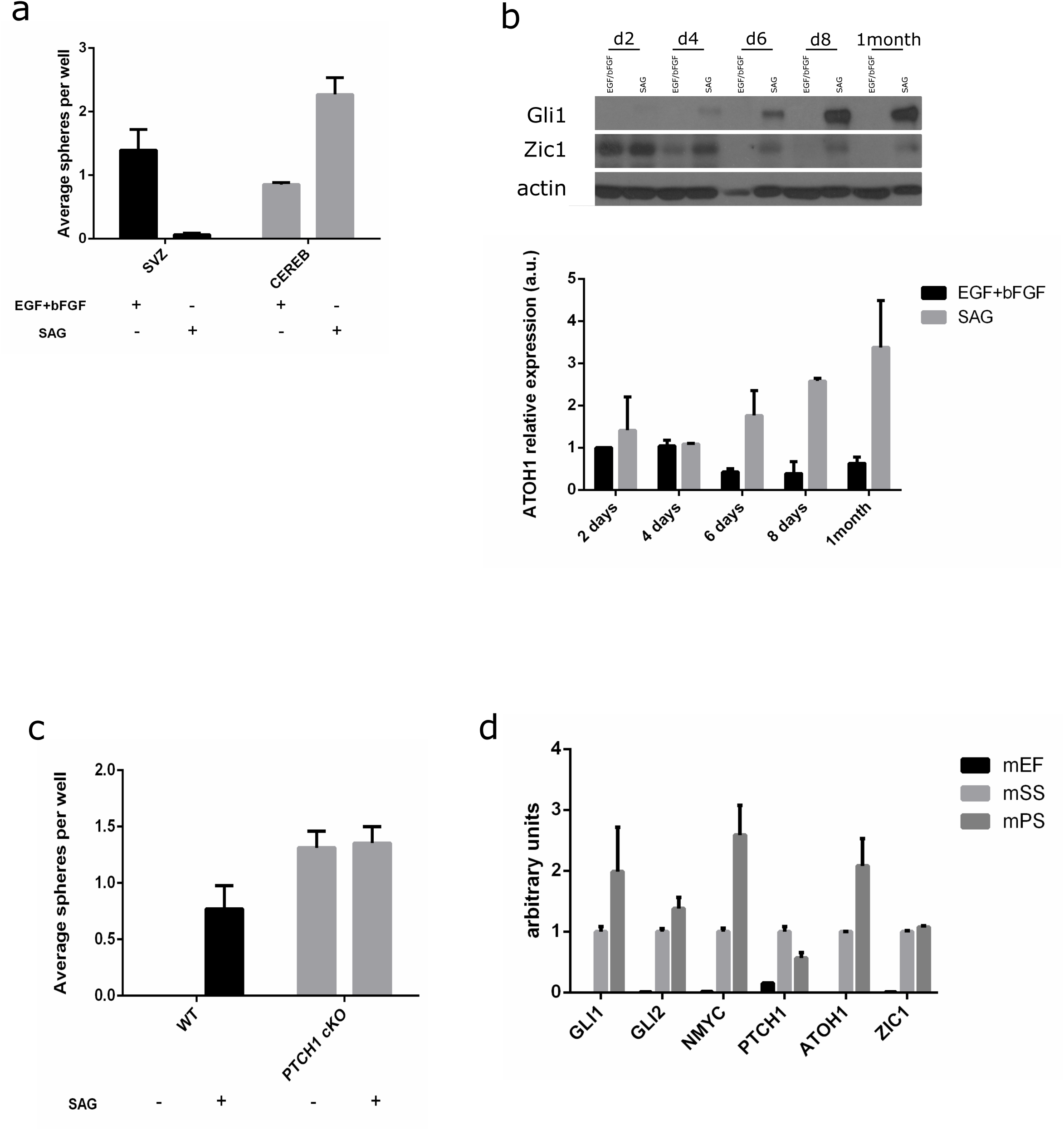
mSS cells belong to the GCP neurogenic lineage. a) Clonogenic assay carried out using explant cells from different sources (SVZ and cerebellum) and exposed to different treatments (SAG and EGF/bFGF). b) Western blot showing GLI1 and ZIC1 protein levels (top) and QPCR to assay ATOH1 levels (bottom) in EGF/bFGF and SAG exposed cultures throughout the duration of 1 month in culture. c) Clonogenic assay carried out with explants from either wild type (WT) or PTCH1 conditional knockout cerebella. d) Comparison of gene expression by QPCR between mEF, mSS and mPS cells.

As proliferating GCP cells are an abundant cell type in the post-natal murine cerebellum I established cultures under EF conditions, as well as SAG treatment and monitored the expression of ZIC1 protein and *atoh1* transcript during the course of the initial culture, as these are both reported as being markers of the GCP lineage. I observe that ZIC1 protein levels gradually decrease in the presence of EF conditions, being undetectable after 6 days, while ZIC1 levels remain detectable after 1 month, concomitant with increased levels of GLI1 (**Fig. 3b**, **top**). Concordantly, ATOH1 transcript levels were gradually enriched in SAG exposed cells while they gradually decreased under EF conditions (**Fig. 3b**, **bottom**). The gradual increase in GCP lineage markers, of which *atoh1* and *zic1* are not transcriptionally activated by the SHH pathway, in SAG exposed cells suggests enrichment of GCP cells at the expense of SAG non-responsive cells.

To further confirm that SAG responsive cells are of the GCP lineage, I investigated if SAG treatment could be substituted with genetic activation of the SHH signaling pathway in the GCP lineage. In ATOH1-CRE; PTCH1 fl/fl mice CRE recombinase expression is driven in cells of the GCP lineage (Yang et al., 2008). Clonogenic assay was carried out on cells originating from p7 cerebella of either ATOH1-CRE; PTCH1 fl/fl mice or WT littermates and clonogenicity was observed with SAG treatment, while only conditional KO cells gave rise to spheres in the absence of SAG (**Fig. 3c**), which will be referred to as murine PTCH KO derived spheres (mPS). Therefore, I maintain, that cells of the GCP lineage respond to SAG treatment with SHH signaling pathway activation and proliferation under these conditions. To further establish the identity of the GCP derived cell lines with respect to mEF neurospheres I assayed expression of key SHH pathway components (*gli1*, *gli2*, *nmyc*, *ptch1*) and GCP lineage markers (*atoh1*, *zic1*). All these markers were expressed similarly in mSS and mPS cells, while expression was very low in mEF cells (**Fig. 3d**).

### mSS cells continuously require SHH pathway stimuation

To assay the requirement of mSS cells for exogenous agonist of the SHH signaling pathway I challenged mSS cells, which had been in culture for over 10 weeks, by removing SAG from the culture medium. I observed a marked decrease in clonogenic capability (**Fig. 4a**).

**Figure 4.**
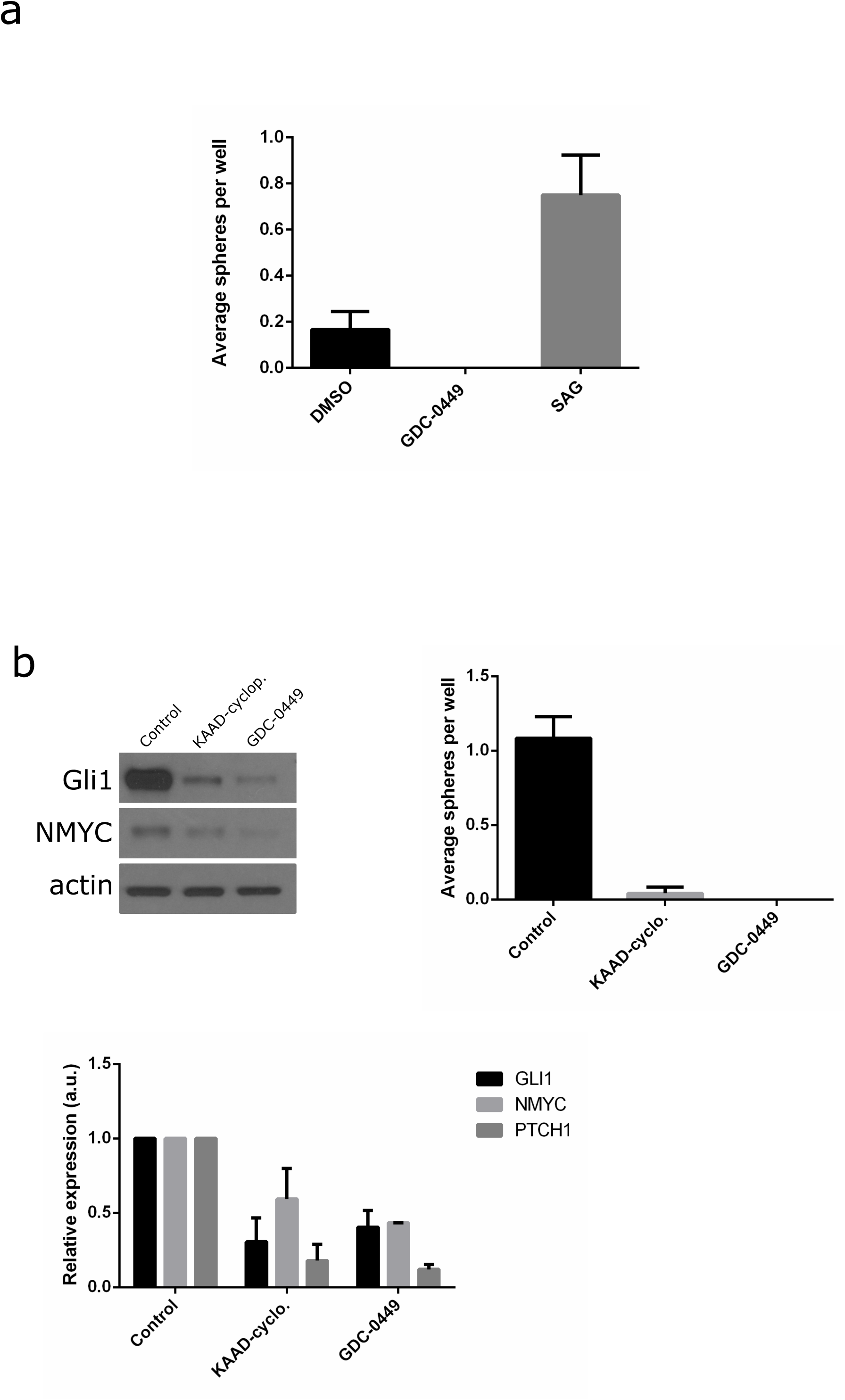
mSS cells maintain dependence on SHH pathway after long-term culturing. a)Clonogenic assay on long-term cultured mSS cells upon derivation from SAG supplementation. b) Western blot and QPCR assaying SHH pathway members (left-top, bottom) and clonogenic assay (right) in Smoothened inhibitor treated mSS cells.

To further confirm the continuous requirement for Smoothened stimulation for SHH pathway activation and proliferation of mSS cells, I treated these cells with either KAAD-cyclopamine or GDC-0449, which are inhibitors of the Smoothened protein. In agreement with the above observations, clonogenicity diminishes with concomitant downregulation of SHH pathway components in terms or transcript and protein levels (**Fig. 4b**).

### mSS cells maintain neural lineage commitment after long-term culture

It has been reported that the plasticity of neural progenitors can be modified after long periods of time in cell culture (Conti and Cattaneo, 2010). To assay the capabilities of mSS cells to differentiate *in vitro* I exposed paired mSS and mEF neurospheres to medium containing serum, a condition reported to promote neurosphere differentiation towards the astrocytic lineage. Both cell types attach to the dish within 24 hours and after 5 days in these conditions mSS cells acquire a bipolar neuronal morphology while mEF neurospheres acquire glial morphology (**Fig. 5a**).

**Figure 5.**
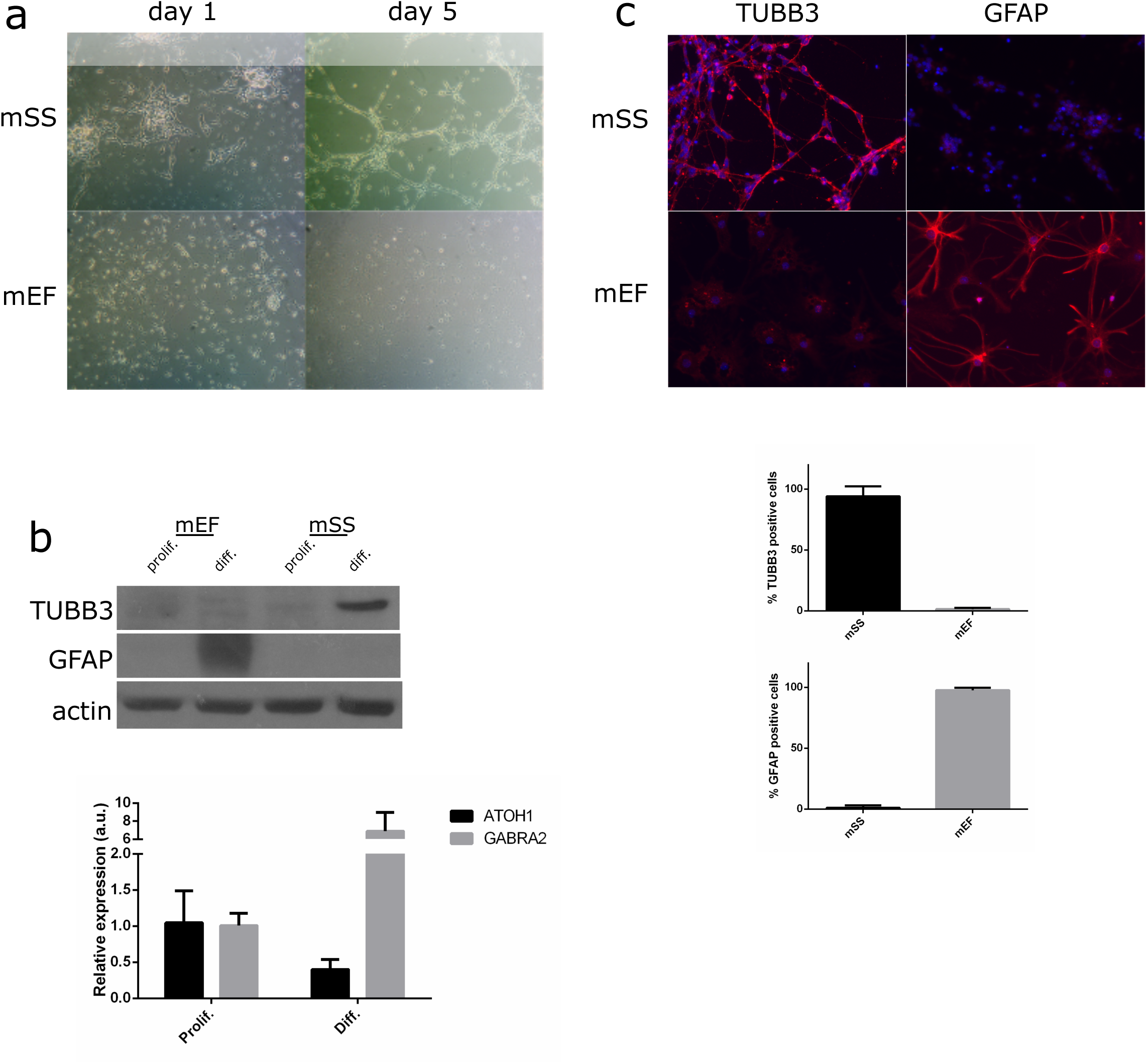
mSS cells are neurogenic even after long-term culture. a) Contrast photo of mSS and mEF cells at 1 day or 5 days in differentiation conditions. b) top) Western blot comparing beta3-tubulin(TUBB3) or GFAP levels between 5 day differentiated mSS or mEF cells. bottom) QPCR of ATOH1 and GABRA2 in differentiated mSS cells. c) Immunofluorescence for TUBB3 or GFAP in mSS or mEF cells at 5 days in differentiation conditions. Histograms (bottom) express relative numbers of each cell type.

To characterize the fate of differentiated neurospheres more precisely I assayed the protein levels of beta3-tubulin and GFAP, markers of neuronal and astrocytic differentiation, respectively. In agreement with the above mentioned morphological changes undergone in differentiation medium mSS cells up-regulated expression of beta3-tubulin, on the other hand mEF neurospheres expressed high levels of GFAP under these conditions (**Fig. 5b**, **top**).

After having established neuronal lineage commitment for mSS cells I assayed expression of *gabra2* and *atoh1* in differentiated cells. In accordance with the GCP identity of mSS cells, *atoh1* transcript levels decreased, while *gabra2* levels accumulated upon differentiation (**Fig. 5b**, **bottom**).

I also analyzed differentiated mEF and mSS cell cultures by immunofluorescence for beta3-tubulin and GFAP. In accord with the mutual exclusivity of GFAP and beta3-tubulin between the two differentiated cell pools, mEFS cells gave rise to a homogeneous population of GFAP positive cells while mSS cells gave rise to a significant majority of beta3-tubulin staining cells (**Fig. 5c**).

In summary, I start out by demonstrating generation of novel MB tumor spheres from murine conditional transgenic tumors. I next explore if the principles of neurosphere culture generation can be extended to non-transformed GCP lineage cells. All in all I find that under favorable culture conditions, GCPs give rise to neurosphere cultures that proliferate extensively, continuously depend on SHH pathway stimulation and maintain commitment and differentiation potential towards their original neural lineage.

## Discussion

I report the generation of cell lines which proliferate in dependence of constant activation of the SHH signaling pathway, from p7 murine cerebellar explants. These cells, termed mSS, have characteristics of GCP cells, such as expression of *atoh1* and *zic1*, as well as of previously reported neural progenitor cell cultures, such as expression of *sox2* and *nestin*. Further, mSS cells maintain extensive cell proliferation. mSS cells therefore combine hallmark properties of both the transient GCP derived *in vitro* systems and conventional neurosphere cultures.

The unlimited proliferation capability of mSS cells were unexpected because GCP cell cultures were previously defined as being transient even in the presence of mitogenic stimuli, such as SHH or the pharmacological smoothened agonist SAG (Miyazawa et al., 2000; Wechsler-Reya and Scott, 1999). On the other hand, extensive self-renewal of TA cells cultured in neurosphere conditions has been reported also in other contexts of neurogenesis (Doetsch et al., 2002).

As was the case in the context of SHH subtype MB, the choice of cell culture medium was key in maintaining activity of the SHH pathway. Cerebellum derived neurospheres cultured in the presence of EGF and bFGF lose activity of the SHH pathway, which is consistent with previous observations. The fact that mono-layer GCP cultures are limited in their expansion could be due to contact between the cell and the plastic substrate, or serum treatment, which is often included in these protocols.

In the past, MB and SHH research have relied on adhesive cell cultures, such as human tumor derived cell lines or murine 3T3 cells (Götschel et al., 2013; Taipale et al., 2000). It is significant to note, that these cell lines require hyperconfluecy and low serum conditions before they become receptive to exogenous treatments that activate the SHH pathway. These treatment conditions are not the ideal for cancer relevant experiments, which aim to measure proliferation or invasive capability. Indeed, in this work, in addition to confirming that bFGF will inhibit SHH activity in established MB cell lines and I also make the novel observation that serum treatment also has this effect. This could explain the previously made observation that culturing murine MB cells in DMEM + 10% FBS leads to cell lines that do not express the SHH pathway (Sasai et al., 2006). Literature suggests that the SHH inducibility through manipulation of confluency and serum may be mediated by the YAP/TAZ signaling pathway (Tariki et al., 2014).

The generation of mSS cell lines is significant also due to pairing of such lines with tumor derived cell lines, as it allows for assaying properties that are specific to transformed cells, allowing for discovery of therapeutics targeted specifically to cancer cells. Current treatments for medulloblastoma often leave patients neurologically impaired (Palmer et al., 2001, 2007), therefore it is a priority to optimize therapeutic index for treatments to this disease. An attractive experimental design in this spirit would be pairings of conditional knockout cell lines as used in this work. ATOH1-CRE; PTCH1 fl/fl cell lines could be generated from early postnatal cerebella as well as established tumors, further studies can then be carried out under homogeneous experimental conditions, as neither of these cell lines require additional culture supplementation to proliferate.

The adherence to the physiological lineage of mSS cells after even long periods in culture is notable, because in the context of other types of neurosphere culture extensive proliferation has been shown to deplete neurogenic potential of neural derived progenitor cultures (Conti and Cattaneo, 2010). mSS cells therefore distinguish themselves from other kinds of neurosphere lines in that they stably adhere to the neural lineage.

The fact that mSS cultures derive from GCP cells strongly suggests that the cultures originate from TA cells which are suspended in the proliferative state through persistent activation of the SHH pathway. The concept that TA cells of neural origin can be easily cultured for long periods of time has also been observed in cells originating from the murine SVZ niche (Doetsch et al., 2002). In that case, TA cells maintained proliferation in a EGF receptor dependent manner, again highlighting the effect that sustained mitogenic signaling can have on TA cells. Cellular lineages consist of quiescent stem cells and differentiated cells which carry out tissue function. Between these two states, TA cells undergo rapid expansion and therefore may constitute a vulnerability in terms of oncogenic transformation. In the case of the GCP lineage this seems partially to be true because oncogenic mutations even in stem cells manifest only after commitment to the GCP lineage (Schüller et al., 2008; Yang et al., 2008).

### Conclusion

To conclude, in this work I describe how opportune defined cell culture conditions can favor establishment of neurosphere cell lines that adhere to the cerebellar GCP lineage. An important property of these lines is their dependency on mitogenic SHH signaling. Strikingly, neurospheres from normal p7 cerebella proliferate indefinitely in culture, which is unexpected given their normal developmental role as TA cells. Further, these cells, named mSS, remain dependent on exogenous mitogenic stimulus and remain adherent to the GCP lineage when placed under differentiation conditions. Together with their tumor derived counterparts, mSS cells establish an *in vitro* paradigm for the study of GCP lineage cells under diseased and normal conditions. This paradigm is unique as it respects the requirement of this lineage for mitogenic SHH signaling.

## Acknowledgements

I thank Dr. Fredrik Swartling for having generously provided GTML samples and cells.

This work was carried out as part of the Molecular Medicine Ph.D. program at the University of Rome “La Sapienza”. Complete funding information can be found at the following URL: https://web.uniroma1.it/phdmolmed/research-funding/research-funding

The complete Ph.D. thesis can be found archived at the German National Library at the following URL: http://d-nb.info/1120340926

## Bibliography

Ahlfeld, J., Favaro, R., Pagella, P., Kretzschmar, H.A., Nicolis, S., and Schüller, U. (2013). Sox2 requirement in sonic hedgehog-associated medulloblastoma. Cancer Res. 73, 3796–3807.

Browd, S.R., Kenney, A.M., Gottfried, O.N., Yoon, J.W., Walterhouse, D., Pedone, C.A., and Fults, D.W. (2006). N-myc can substitute for insulin-like growth factor signaling in a mouse model of sonic hedgehog-induced medulloblastoma. Cancer Res. 66, 2666–2672.

Butts, T., Green, M.J., and Wingate, R.J.T. (2014). Development of the cerebellum: simple steps to make a “little brain.” Development 141, 4031–4041.

Capdevila, C., Rodríguez Vázquez, L., and Martí, J. (2016). Glioblastoma Multiforme and Adult Neurogenesis in the Ventricular-Subventricular Zone: A Review. J. Cell. Physiol.

Conti, L., and Cattaneo, E. (2010). Neural stem cell systems: physiological players or in vitro entities? Nat. Rev. Neurosci. 11, 176–187.

Doetsch, F., Petreanu, L., Caille, I., Garcia-Verdugo, J.M., and Alvarez-Buylla, A. (2002). EGF converts transit-amplifying neurogenic precursors in the adult brain into multipotent stem cells. Neuron 36, 1021–1034.

Emmenegger, B.A., Hwang, E.I., Moore, C., Markant, S.L., Brun, S.N., Dutton, J.W., Read, T.-A., Fogarty, M.P., Singh, A.R., Durden, D.L., et al. (2013). Distinct roles for fibroblast growth factor signaling in cerebellar development and medulloblastoma. Oncogene 32, 4181–4188.

Gonçalves, J.T., Schafer, S.T., and Gage, F.H. (2016). Adult Neurogenesis in the Hippocampus: From Stem Cells to Behavior. Cell 167, 897–914.

Götschel, F., Berg, D., Gruber, W., Bender, C., Eberl, M., Friedel, M., Sonntag, J., Rüngeler, E., Hache, H., Wierling, C., et al. (2013). Synergism between Hedgehog-GLI and EGFR signaling in Hedgehog-responsive human medulloblastoma cells induces downregulation of canonical Hedgehog-target genes and stabilized expression of GLI1. PloS One 8, e65403.

Kessler, J.D., Hasegawa, H., Brun, S.N., Emmenegger, B.A., Yang, Z.-J., Dutton, J.W., Wang, F., and Wechsler-Reya, R.J. (2009). N-myc alters the fate of preneoplastic cells in a mouse model of medulloblastoma. Genes Dev. 23, 157–170.

Lee, A., Kessler, J.D., Read, T.-A., Kaiser, C., Corbeil, D., Huttner, W.B., Johnson, J.E., and Wechsler-Reya, R.J. (2005). Isolation of neural stem cells from the postnatal cerebellum. Nat. Neurosci. 8, 723–729.

Miyazawa, K., Himi, T., Garcia, V., Yamagishi, H., Sato, S., and Ishizaki, Y. (2000). A role for p27/Kip1 in the control of cerebellar granule cell precursor proliferation. J. Neurosci. Off. J. Soc. Neurosci. 20, 5756–5763.

Northcott, P.A., Jones, D.T.W., Kool, M., Robinson, G.W., Gilbertson, R.J., Cho, Y.-J., Pomeroy, S.L., Korshunov, A., Lichter, P., Taylor, M.D., et al. (2012). Medulloblastomics: the end of the beginning. Nat. Rev. Cancer 12, 818–834.

Palmer, S.L., Goloubeva, O., Reddick, W.E., Glass, J.O., Gajjar, A., Kun, L., Merchant, T.E., and Mulhern, R.K. (2001). Patterns of Intellectual Development Among Survivors of Pediatric Medulloblastoma: A Longitudinal Analysis. J. Clin. Oncol. 19, 2302–2308.

Palmer, S.L., Reddick, W.E., and Gajjar, A. (2007). Understanding the Cognitive Impact on Children Who are Treated for Medulloblastoma. J. Pediatr. Psychol. 32, 1040–1049.

Pastrana, E., Silva-Vargas, V., and Doetsch, F. (2011). Eyes wide open: a critical review of sphere-formation as an assay for stem cells. Cell Stem Cell 8, 486–498.

Rahman, M., Reyner, K., Deleyrolle, L., Millette, S., Azari, H., Day, B.W., Stringer, B.W., Boyd, A.W., Johns, T.G., Blot, V., et al. (2015). Neurosphere and adherent culture conditions are equivalent for malignant glioma stem cell lines. Anat. Cell Biol. 48, 25–35.

Reynolds, B.A., and Weiss, S. (1992). Generation of neurons and astrocytes from isolated cells of the adult mammalian central nervous system. Science 255, 1707–1710.

Rios, I., Alvarez-Rodríguez, R., Martí, E., and Pons, S. (2004). Bmp2 antagonizes sonic hedgehog-mediated proliferation of cerebellar granule neurones through Smad5 signalling. Dev. Camb. Engl. 131, 3159–3168.

Sasai, K., Romer, J.T., Lee, Y., Finkelstein, D., Fuller, C., McKinnon, P.J., and Curran, T. (2006). Shh pathway activity is down-regulated in cultured medulloblastoma cells: implications for preclinical studies. Cancer Res. 66, 4215–4222.

Schüller, U., Heine, V.M., Mao, J., Kho, A.T., Dillon, A.K., Han, Y.-G., Huillard, E., Sun, T., Ligon, A.H., Qian, Y., et al. (2008). Acquisition of granule neuron precursor identity is a critical determinant of progenitor cell competence to form Shh-induced medulloblastoma. Cancer Cell 14, 123–134.

Taipale, J., Chen, J.K., Cooper, M.K., Wang, B., Mann, R.K., Milenkovic, L., Scott, M.P., and Beachy, P.A. (2000). Effects of oncogenic mutations in Smoothened and Patched can be reversed by cyclopamine. Nature 406, 1005–1009.

Tariki, M., Dhanyamraju, P.K., Fendrich, V., Borggrefe, T., Feldmann, G., and Lauth, M. (2014). The Yes-associated protein controls the cell density regulation of Hedgehog signaling. Oncogenesis 3, e112.

Vanner, R.J., Remke, M., Gallo, M., Selvadurai, H.J., Coutinho, F., Lee, L., Kushida, M., Head, R., Morrissy, S., Zhu, X., et al. (2014). Quiescent sox2(+) cells drive hierarchical growth and relapse in sonic hedgehog subgroup medulloblastoma. Cancer Cell 26, 33–47.

Vescovi, A.L., Galli, R., and Reynolds, B.A. (2006). Brain tumour stem cells. Nat. Rev. Cancer 6, 425–436.

Wechsler-Reya, R.J., and Scott, M.P. (1999). Control of neuronal precursor proliferation in the cerebellum by Sonic Hedgehog. Neuron 22, 103–114.

Yang, Z.-J., Ellis, T., Markant, S.L., Read, T.-A., Kessler, J.D., Bourboulas, M., Schüller, U., Machold, R., Fishell, G., Rowitch, D.H., et al. (2008). Medulloblastoma can be initiated by deletion of Patched in lineage-restricted progenitors or stem cells. Cancer Cell 14, 135–145.

Zhao, X., Ponomaryov, T., Ornell, K.J., Zhou, P., Dabral, S.K., Pak, E., Li, W., Atwood, S.X., Whitson, R.J., Chang, A.L.S., et al. (2015). RAS/MAPK Activation Drives Resistance to Smo Inhibition, Metastasis, and Tumor Evolution in Shh Pathway-Dependent Tumors. Cancer Res. 75, 3623–3635.

